# On Levodopa interactions with brain disease proteins at the nanoscale

**DOI:** 10.1101/2024.11.15.623204

**Authors:** Talia Bergaglio, Nico Kummer, Shayon Bhattacharya, Damien Thompson, Silvia Campioni, Peter Niraj Nirmalraj

## Abstract

The cerebral accumulation of α-Synuclein (α-Syn) and amyloid β-1-42 (Aβ-42) proteins are known to play a crucial role in the pathology of neurocognitive disorders such as Parkinson’s disease (PD). Currently, Levodopa (L-dopa) is the dopamine replacement therapy for treating bradykinetic symptoms visible in PD patients. Here, we use atomic force microscopy to evidence at nanometer length scales the effects of L-dopa on the morphology of α-Syn and Aβ-42 protein fibrils. L-dopa treatment reduces the length and diameter of both types of protein fibrils, with a stark reduction observed for Aβ-42 both in physiological buffer and human spinal fluid. The insights gained on Aβ-42 fibril disassembly from the nanoscale imaging experiments are substantiated using atomic-scale molecular dynamics simulations. Our results reveal the mechanism governing L-dopa-driven reversal of protein aggregation, which may be useful in drug design of small molecule drugs for potentially treating neurocognitive disorders and provide leads for designing chemical effector-mediated disassembly of protein architectures.

---

Alzheimer’s and Parkinson’s disease are characterized by the progressive accumulation of protein aggregates in brain synapses^1^. Yet, there is a stark difference in the chemical structure of the proteins implicated in each neurodegenerative disorder. In Parkinson’s disease (PD), aggregation of the misfolded *α*-synuclein (*α*-Syn) protein^2-4^, from its native soluble disordered state to β-sheet-rich insoluble fibril structures, drives the formation of toxic intracellular aggregates known as Lewy bodies and Lewy neurites. This leads to the degeneration of dopaminergic neurons within the substantia nigra, switching off synthesis of the neurotransmitter dopamine in the midbrain basal ganglia structure^5-7^. Conversely, in Alzheimer’s disease (AD), the aggregation of amyloid β peptide (chiefly, Aβ-42)^8, 9^ and tau protein^10^ in the brain tissue results in the formation of amyloid plaques and neurofibrillary tangles, respectively, triggering neurocognitive impairments^11^. There is emerging evidence that individuals with PD frequently exhibit non-motor symptoms typical of AD patients, with *α*-Syn and Aβ accumulations co-detected in the brain of PD and AD patients^12, 13^. Importantly, pathogenic processes triggering the onset of AD and PD typically occur several decades before the emergence of visible signs of memory and cognitive deficits^14^, indicating that the gradual changes in *α*-Syn and Aβ protein structure from monomers, oligomers, and protofibrils to fibrils progressively disrupt neuronal function and connectivity^15^. These pathological protein aggregates are present in both blood^16^ and cerebrospinal fluid (CSF)^17^ of PD and AD patients, which provides an opportunity to monitor disease progression before the onset of clinical symptoms through development of fluid biomarkers aimed at the detection and quantification of α-Syn and Aβ-42 proteins^18^.

Levodopa (L-dopa) is a first-line drug in the pharmacological management of PD. As a precursor to dopamine, L-dopa replenishes dopamine levels in the brain’s depleted regions, providing effective relief from motor symptoms in PD patients^19^. While the distribution and degree of *α*-Syn buildup in different regions of the brain of individuals with PD are well-documented, the effect of long-term dopamine replacement therapy on *α*-Syn aggregation remains unknown^20^. Importantly, prolonged L-dopa therapy can be associated with motor fluctuations, dyskinesia, and a consequent reduction in the effectiveness of a given L-dopa dose^21^. The generation of free radicals, as well as L-dopa-induced toxicity to dopaminergic neurons, may, in turn, accelerate the neurodegenerative processes underlying PD^22^. Although L-dopa cannot arrest or slow down the advancement of PD nor reverse the course of the disease, *in vitro* studies have demonstrated that L-dopa can effectively hinder the formation of *α*-Syn fibrils and promote the disassembly of pre-existing fibrils^23, 24^. The effect of L-dopa on synapses is schematically shown in Fig. 1A, comparing healthy, diseased, and L-dopa-treated neuronal interfaces. While L-dopa is primarily used in the management of PD, the effects of the dopamine precursor on Aβ pathology have also been investigated due to the synergy of PD and AD pathology^25^. Specifically, *in vitro* studies using Thioflavin T (ThT) fluorescence have shown that L-dopa may alter the morphology of Aβ aggregates by inhibiting Aβ fibrillation^23, 24^. While the investigation into the therapeutic efficacy of L-dopa on cognitive decline in dementia remains ongoing^26, 27^, the substantial involvement of dopamine in learning and memory consolidation has sparked considerable interest in its potential application beyond PD^28, 29^, including mitigation of AD-related pathology by L-dopa targeting of Aβ aggregation^30^. Hence, understanding the complex mechanisms governing *α*-Syn and Aβ aggregation and the mode of action of dopamine replacement therapies is essential for effective therapeutic interventions designed to halt PD progression and, specifically, PD-related dementia. Although previous studies using Thioflavin T (ThT), kinetics binding assays have suggested a significant reduction in *α*-Syn and Aβ aggregation in the presence of L-dopa^23, 24^, nanoscale characterization techniques, such as atomic force microscopy (AFM), can provide high-resolution visualization and quantitative measurement of the mechanical properties, aggregation patterns, and morphological changes induced by L-dopa^23, 31, 32^. Molecular insights into protein–drug interactions are crucial for predicting the clinical outcomes of L-dopa treatment in PD and AD patients. Previously, we resolved and quantified the size, shape, and morphology of diverse protein aggregates formed along the primary aggregation pathway of wild-type *α*-Syn and Aβ-40 and Aβ-42 *in vitro* using atomic force microscopy (AFM)^31, 33^

**Figure 1:**
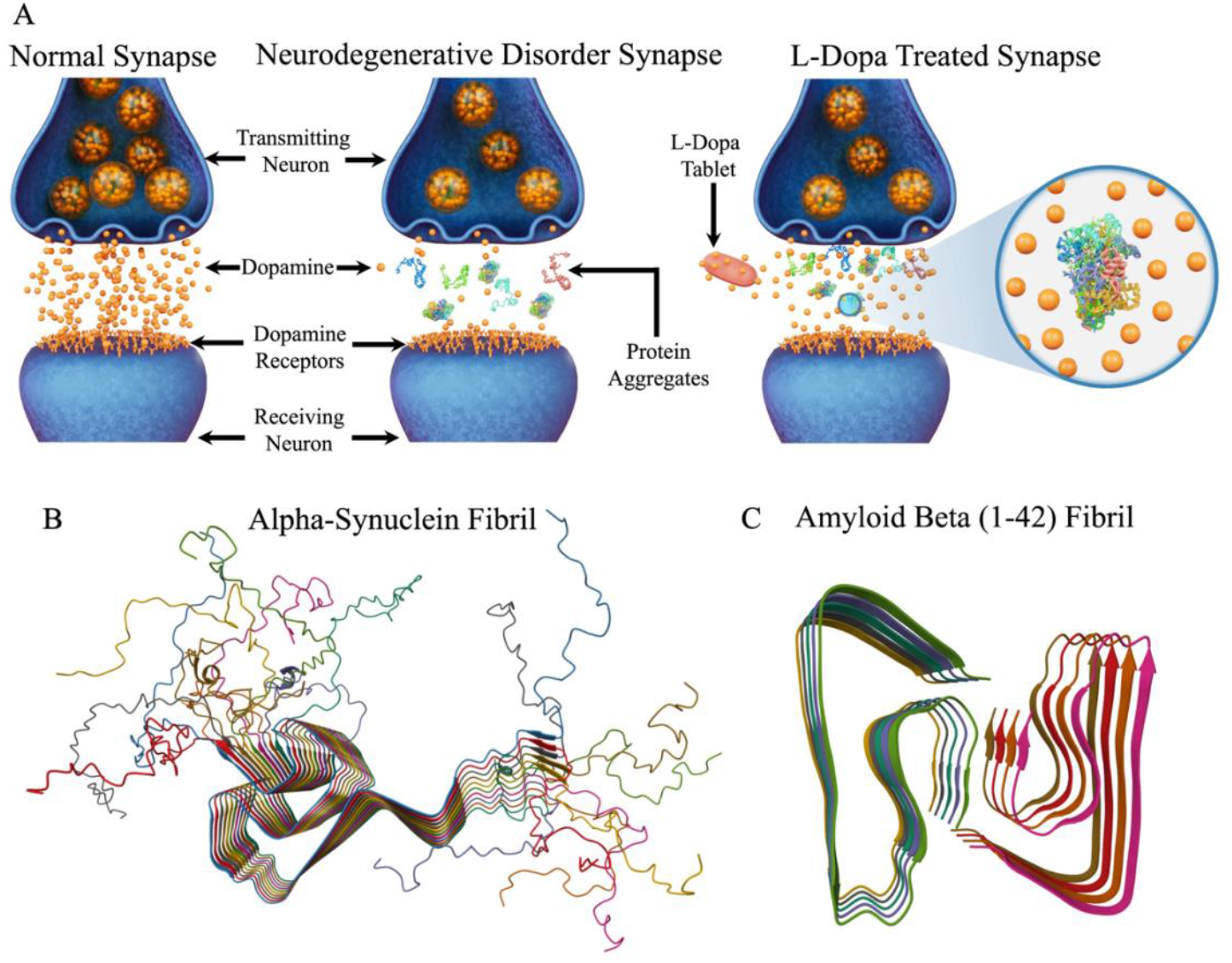
Study rationale and pathological protein structure. (A) Schematic detailing the differences between normal synapse, neurocognitive disorder synapse, and levodopa-treated synapse. The objects shown are not to scale. Understanding the impact of levodopa on protein aggregates such as alpha-synuclein (B, protein data bank identifier: 2NOA) and amyloid beta (1-42) fibril (C, protein data bank identifier: 5OQV) implicated in neurodegenerative disorders such as Parkinson’s and Alzheimer’s disease using nanoscale imaging is the focus of the present study.

In the present work, we extend the application of AFM to investigate the effects of L-dopa on *α*-Syn and Aβ-42 fibrils at nanometer length scales. The atomically resolved structures of *α*-Syn and Aβ-42 fibrils used as models in our previous works^31, 33^ are shown in Figures 1B and 1C, respectively. When treated with 100 μM L-dopa, a reduction in the length and diameter of fibrils was observed for *α*-Syn and Aβ-42 (see Supporting Information section for protein solution preparation). Control studies in human CSF revealed a significant decrease in fibril length and diameter when compared to untreated Aβ-42-CSF samples, indicative of protein fibrillar disassembly through the action of L-dopa. Molecular dynamics simulations revealed the formation of a physisorbed layer of L-dopa on the fibrils. This layer facilitates their destabilization and masking of the aggregation-prone hydrophobic cores of the released low molecular weight oligomers, thus inhibiting their reincorporation into the fibrils.

We first performed AFM characterization of *α*-Syn fibrils that were incubated with and without 100 μM L-dopa for six days under mechanical agitation at 37°C. Figure 2a shows an AFM height image recorded after depositing the untreated *α*-Syn solution, showing aggregated *α*-Syn fibrils. The AFM height image for the L-dopa treated sample (Figure 2B) shows the presence of *α*-Syn fibrils and L-dopa particles (white arrows, visible on the fibrils and throughout the sample). To assess if the detected spherical particles in Fig. 2B could represent L-dopa particles, we calculated the mean size of all the spherical particles resolved in the AFM images where *α*-Syn proteins were incubated together with L-dopa. Based on single particle size analysis, we calculated a mean L-dopa particle size of 10.5 ± 1.95 nm (Fig. 2C). Such spherical particle sizes were not detected when *α*-Syn proteins were incubated in the absence of L-dopa, suggesting that spherical particles are L-dopa molecules present as aggregates. To quantify the effect of L-dopa treatment on *α*-Syn aggregation, we extracted the height and length distribution, as well as the persistence length measurement, of *α*-Syn fibrils incubated without and with 100 μM L-dopa. A significant decrease in fibril length (Figure 2D) and height (Figure 2E) was observed for the α-Syn fibrils incubated with L-dopa. A two-sample t-test revealed a significant reduction in fibril length (Figure 2D) when incubated with 100 μM L-dopa (610 ± 330 nm) compared to the untreated fibrils (860 ± 590 nm). Similarly, the mean height (Figure 2E) of the individual fibrils was also reduced upon treatment with L-dopa (5.20 ± 1.8 nm) compared to the control condition (5.96 ± 1.95 nm).

**Figure 2:**
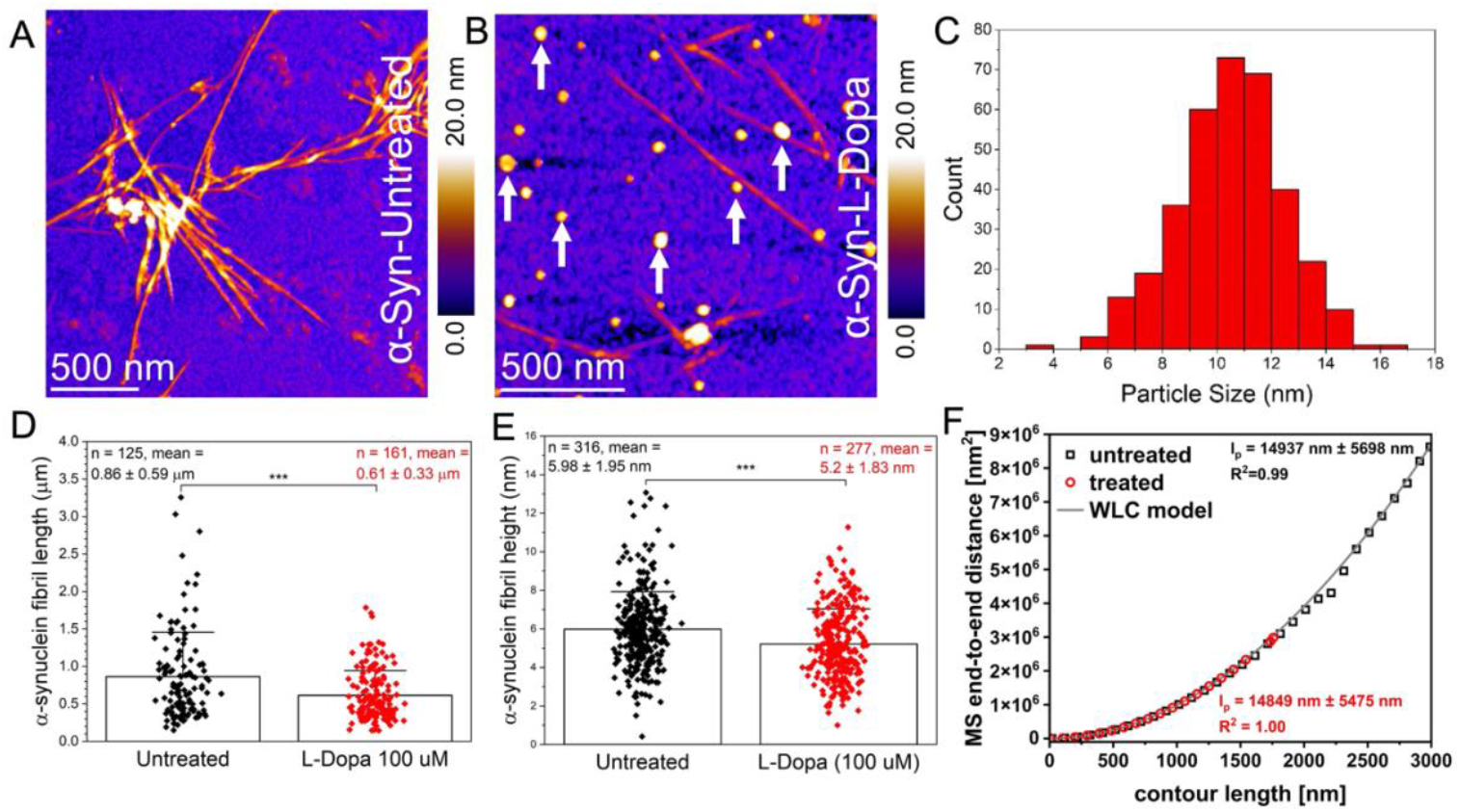
Nanoscale imaging of untreated and L-dopa treated α-Syn in buffer salt solution. (A) AFM height map of untreated α-Syn protein fibrillar aggregates deposited on gold surface. (B) AFM height image of α-Syn protein monomers incubated with 100 μM L-dopa and deposited on a gold surface. (C) Size distribution of spherical particles of 10.53 ± 1.96 nm, based on AFM height data. (D-E) Plot of the mean α-Syn fibril length and height values obtained in untreated samples (coded in black) and samples incubated with L-dopa (coded in red). Error bars indicate the standard deviation from the mean. (F) The persistence length of untreated α-Syn fibrils (hollow sphere trace) and L-dopa treated (red trace) α-Syn fibrils. The worm-like chain model (WLC) is plotted as the dark gray line.

Additionally, we assessed the nanomechanical properties of the α-Syn fibrils with and without L-dopa treatment. Figure 2F shows the mean square end-to-end distance as a function of the contour length for untreated fibrils (blue) and treated fibrils (red). The calculated persistence length suggested a very small difference in mechanical properties between experimental conditions, where untreated *α*-Syn fibrils showed a slightly higher average persistence length (14.94 ± 5.70 μm) than L-dopa-treated α-Syn fibrils (14.85 ± 5.48 μm). Previously, we studied the effect of ibuprofen on blood and observed a concentration and time-dependent effect on the morphology of red blood cells (RBCs), characterized by the formation of spicules on the RBC membrane and the transition from normocytes into echinocytes^34^. To evaluate the effect of L-dopa on blood, we conducted similar measurements, as detailed in Fig. S1; however, we did not detect any L-dopa-induced structural changes in RBC morphology. Our findings indicate that, while L-dopa plays a role in the disassembly of *α*-Syn protein aggregation, there were no corresponding structural alterations in RBC morphology, suggesting that L-dopa likely has minimal deleterious hematological effects upon interaction with blood.

Next, we characterized using AFM the Aβ-42 protein aggregates under identical peptide concentration as the α-Syn experiments (see Supporting Information for details on Aβ-42 peptide solution preparation). Note: The incubation temperature was kept at 37°C for both α-Syn and Aβ-42 peptide solution preparation, but the incubation time was different (6 days for α-Syn and 24 hours for Aβ-42). It is known from previous *in-vitro* studies Aβ-42 tends to aggregate faster even along the primary pathway^31^, evidenced by early onset fibril formation when compared to Aβ-40 and α-Syn proteins. We recently showed that Aβ-42 proteins tend to generate oligomers on the surface of primary fibrils through an accelerated secondary nucleation pathway^35^. Based on these previous studies, we implemented a shorter incubation protocol for Aβ-42 compared to α-Syn proteins, as that was sufficient to generate the necessary mature fibrils. The AFM height map of untreated Aβ-42 (Fig. 3A) confirms the predominant presence of mature Aβ-42 fibrils. The presence of mostly fibrils and the absence of smaller oligomeric particles on the solid surface suggests that the 24-hour incubation period results in the saturation phase of Aβ-42 assembly. The L-dopa treated Aβ-42 sample (Fig. 3B) reveals shorter fibril fragments (indicated by black arrows in Fig. 3C), which were not detected for the untreated Aβ-42 proteins, together with an overall reduction of the height of the fibrils. Figure 3D is a high-resolution AFM topograph of an individual L-dopa treated Aβ-42 fibril. The diameter, when measured at multiple points along the elongated fibril (no nodular morphology typical of protofibrils) length, was ∼2 nm.

**Figure 3.**
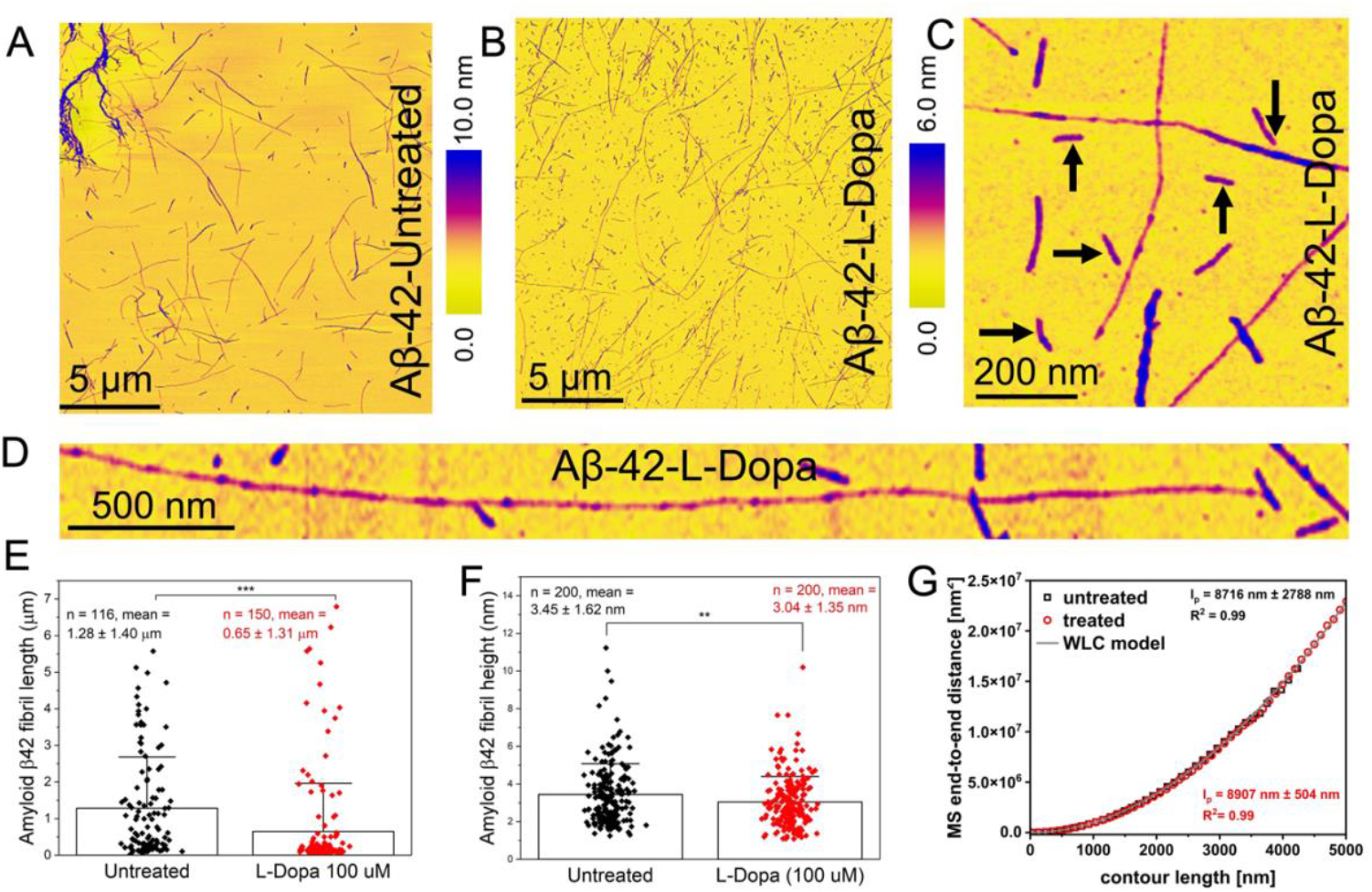
Characterization of untreated and L-dopa treated Aβ-42 in physiological buffer. (A) Large-area AFM image showing untreated Aβ-42 fibrils. (B) AFM image showing L-dopa treated Aβ-42 fibrils of diverse lengths formed after incubating Aβ-42 peptides with L-dopa. (C) AFM image of L-dopa treated Aβ-42 proteins confirming the presence of short fibrils (indicated by black arrows). (D) AFM image of a single Aβ-42 fibril treated with L-dopa. (E) Distribution of fibril length for untreated Aβ-42 (black color-coded) and L-dopa treated Aβ-42 fibrils (red color-coded). (F) Distribution of fibril height for untreated Aβ-42 (black color-coded) and L-dopa treated Aβ-42 fibrils (red color-coded). (G) Combined plot of MS end-to-end distance versus contour length for untreated Aβ-42 fibrils (hollow square trace) and L-dopa treated Aβ-42 fibrils (red sphere trace).

Based on AFM measurements on several such individual fibrils we calculated a mean fibril length of (1.28 ± 1.4 μm) for untreated (n=116, black plot Fig. 3E) and (0.65 ± 1.31 μm) for L-dopa treated Aβ-42 fibrils (n=150, red plot Fig. 3E). Likewise, and in common with the trends observed for α-Syn, the AFM data confirmed an overall reduction in fibril diameter (Fig. 3F) but no measurable differences in persistence length (Fig. 3G) after treatment with L-dopa. Based on the nanoscale imaging experiments conducted on the aggregated forms of *α*-Syn and Aβ-42 proteins in the presence and absence of L-dopa, we observed the most noticeable morphological changes (reduction in fibril length and diameter) to be associated with L-dopa-treated Aβ-42 species. All measurements detailed above were conducted on synthetically prepared proteins in physiological buffer salt solutions, which are much simpler in composition (mainly water, sodium chloride, and phosphate) compared to body fluids such as CSF (water, proteins, ions, and other organic electrolytes). Although information obtained from morphological studies on pathological proteins in buffer salt solutions can provide some insights into protein aggregation mechanisms, it is important to directly study such processes in CSF, as it is a more biologically relevant environment for both therapy and diagnostics.

To test the effect of L-dopa on protein assembly in CSF, we purchased commercially available samples of Aβ-42 proteins enriched in a healthy human CSF from Sigma Aldrich (see materials and methods section in supporting information for additional details). 5 µL of Aβ-42 in the CSF sample was drop-casted on the gold substrate and imaged using AFM, followed by air-drying the sample for five hours under standard laboratory conditions. Figure 4A is an AFM image showing the presence of both large spherical particles and dense fibrils, which we classify as Aβ-42 protein aggregates. The fibrils appear to be closely packed in CSF, as observed in the zoomed-in image (Fig. 4B), which contrasts with the isolated nature of the Aβ-42 fibrils deposited from buffer salt solution on gold substrate. Inspecting the Aβ-42 CSF sample incubated with L-dopa (concentration: 100 μM) for six days at 37°C under mechanical agitation and deposition on a gold surface revealed a reduction in the prevalence, length, and height of the Aβ-42 fibrils. Figure 4C is a representative AFM image of L-dopa treated Aβ-42 protein aggregates in CSF. The spherical particles in CSF were still present after L-dopa incubation (indicated by white arrows). However, densely packed Aβ-42 fibrils previously resolved in untreated CSF samples (Fig. 4A-B) were no longer prevalently detected in L-dopa-treated CSF samples. A small population of fibrils was still observed, but the fibrils were of reduced diameter, as indicated by the yellow arrow in Fig. 4C. The distribution in length and height of untreated Aβ-42 fibrils (black plots) in CSF and L-dopa treated Aβ-42 fibrils in CSF (red plots) is shown in Figs. 4D and E, respectively. The combined height distribution of untreated (black plots) and L-dopa treated (red plots) Aβ-42 protein fibrils in buffer solution and CSF is summarized in Fig. 4F. Taken together, the AFM measurements on Aβ-42 aggregates in CSF confirmed that L-dopa also disassembles Aβ-42 fibrils evidenced through the measured reduction in fibril length and height after L-dopa treatment.

**Figure 4.**
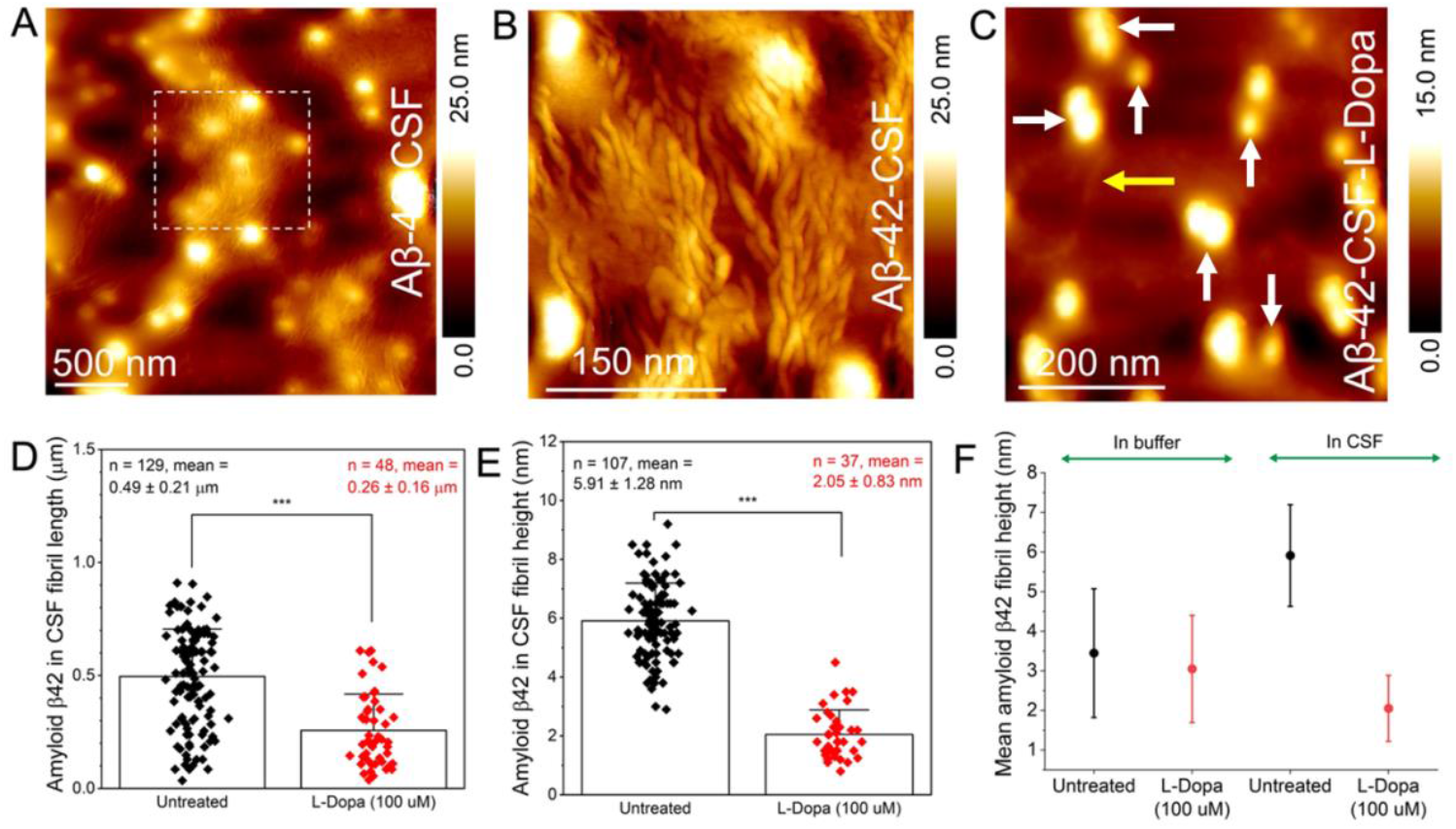
Direct imaging of untreated and L-dopa treated Aβ-42 in human CSF. (A) AFM image revealing the presence of both fibrillar and spherical Aβ-42 aggregates in CSF. (B) High-resolution AFM image of untreated Aβ-42 fibrils resolved within the white dash box in panel A. (C) Large-area AFM image of L-dopa (100 μM) treated Aβ-42 aggregates showing the presence of mostly spherical (indicated by white arrows) and lower prevalence of fibrils (indicated by yellow arrow). (D) Distribution of fibril length for untreated Aβ-42 (black color-coded) and L-dopa treated Aβ-42 fibrils (red color-coded) in CSF. (E) Distribution of fibril height for untreated Aβ-42 (black color-coded) and L-dopa treated Aβ-42 fibrils (red color-coded) in CSF. (F) Mean Aβ-42 fibril height measured in buffer solution and CSF, without (colored black, mean_buffer_: 3.45 ± 1.62 nm; mean_CSF_: 5.91 ± 1.28 nm) and with incubation with100 μM L-dopa (colored red, mean_buffer_: 3.05 ± 1.35 nm; mean_CSF_: 2.05 ± 0.83 nm).

To further understand and quantify the interfacial interactions of L-dopa with the deposited Aβ-42 protein fibrils, we performed atomic-scale molecular dynamics simulations. We sample multiple Aβ-42 fibril fold morphologies by creating starting structures in the LS-shaped fold solved by cryo-EM (PDB code 5OQV^36^) and the double-horseshoe-shaped structure solved by solution NMR (PDB code 2NAO^37^). The protein assembly was placed at the gold substrate in a large water box for simulations without and with 1.3 mM of L-dopa (see Methods section in Supporting Information), and the L-dopa interacted freely with the protein during 0.1 μs of free dynamics (Figs. 5A, Figs. S2C-F). The models reveal a greater loss in native contacts (Q(X))^38^ (Fig. 5B) with the partial unfolding of β-sheet to random coil (Figs. S2E-H) for Aβ-42 treated with L-dopa compared to untreated protofibrils. To account for the fibril thermodynamic stabilities without and with L-dopa, we computed the protein conformational energies by the Generalized Born using Molecular Volume (GBMV) Solvation Energy method implemented in the CHARMM (v40b2) program^39^. While the thermodynamic stability of the L-dopa-treated LS-shaped Aβ-42 fibril fold is at par with the untreated fold (Fig. 5C), the AD-relevant double-horseshoe-shaped fibril fold showed different thermodynamic stability^40^ and is significantly destabilized when treated with L-dopa (Fig. 5C). This NMR-solved double-horseshoe cross-β fibril polymorph was recognized by the antibodies that typically bind to intracellular deposits and senile plaques in the brains of AD patients.

**Figure 5.**
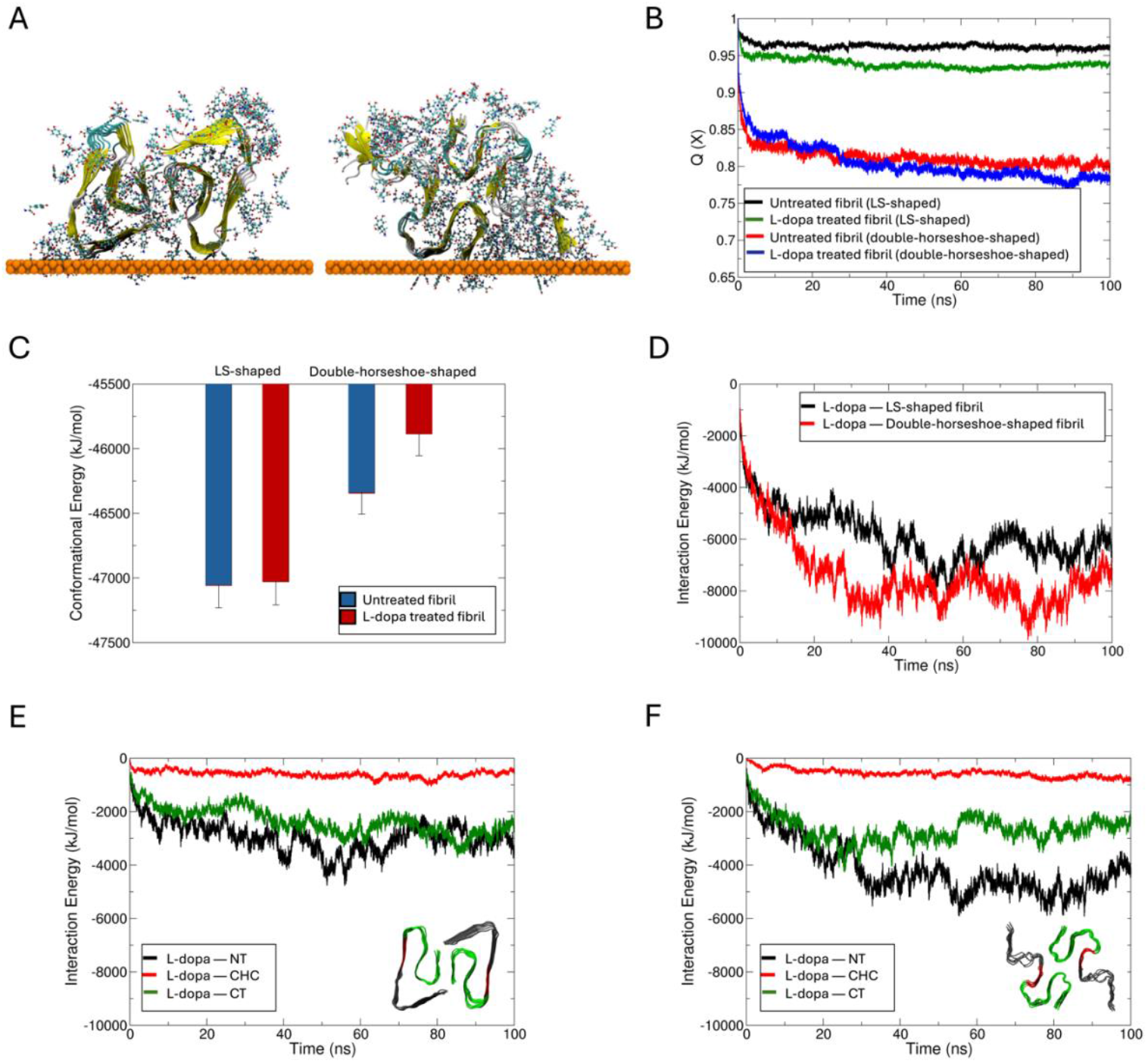
Impact of L-dopa on Aβ-42 protein fibril folds from molecular dynamics simulations. **(A)** The final conformation of Aβ-42 fibril in their LS-shaped fold (PDB code 5OQV^36^) and double-horseshoe-shaped fold (right, PDB code 2NAO^37^) in the presence of L-dopa molecules at the gold–water interface following 100 ns of dynamics. L-dopa molecules within 5 Å of the fibril are shown in ball and stick representation. The protein fibril is shown in its secondary structure representation, and the gold surface is shown as an atomic sphere. Comparison of the time evolution of **(B)** fraction of native contacts (Q(X)) and **(C)** conformational energies between untreated and L-dopa treated fibril folds. Time evolution of interaction energies between L-dopa and **(D)** the two fibril folds, and between L-dopa and the N-terminus (NT, black), central hydrophobic cluster (CHC, red) and C-terminus (CT, green) of **(E)** LS-shaped, and **(F)** double-horseshoe-shaped fibril folds. The two fibril folds with different colored regions are shown as insets.

During the MD simulations, the L-dopa molecules spontaneously formed an ordered cloak around the fibrils driven by Coulombic L-dopa–fibril pairwise interactions (Fig. 5D, Fig. S2G,H). The L-dopa–fibril interactions are majorly contributed by the N-terminus (NT, residues 1-16) of the double-horseshoe-shaped fold followed by the C-terminus (CT, residues 22-42) and the central hydrophobic cluster (CHC, the hydrophobic “hotspots” of aggregation, residues 17-21) (Fig. 5F). By contrast, the L-dopa–NT interactions are not significantly greater than the L-dopa—CT interactions in the LS-fold due to a more structured and less exposed NT than in the double-horseshoe fold (Fig. 5E). The remaining significant L-dopa–CHC interactions suggest that the re-assembly of L-dopa treated etched fibrils may be slowed by blocking the CHC hotspots of aggregation by L-dopa in the released low molecular weight species. Overall, our modeling data strongly supports the finding from AFM maps that the interaction of Aβ-42 fibril with L-dopa leads to destabilization and disassembly of fibrils with high affinity and destabilizing effect of L-dopa molecules around the fibril and favorable L-dopa–CHC contacts may also screen these aggregation sites and prevent reintegration of released oligomers.

In summary, we evidence the disassembly of fibrillar forms of Aβ-42 and α-Syn proteins at the nanoscale upon treatment with L-Dopa. The differences in the size and shape of the individual L-Dopa-treated protein fibrils are recorded in a label-free manner using atomic force microscopy. In particular, the effect of L-Dopa was observed to have a more distinct effect on Aβ-42 fibrils both in physiological solution and in human cerebrospinal fluid. Molecular dynamics simulations quantified the interfacial interactions between L-Dopa and Aβ-42 fibrils. The disassembly of fibrillar structures observed upon L-dopa interaction, particularly with Aβ-42 fibrils, indicates that L-dopa disrupts mature fibrils and may promote the formation of smaller oligomeric species. While the disassembly of fibrillar proteins by L-dopa could represent a therapeutic strategy to halt or reverse disease progression, there is a significant concern that intermediates formed during fibril disassembly, or products of incomplete fibrillation, may exhibit neurotoxic properties^41^. The MD simulations show evidence for the destabilizing effect of L-Dopa with key Aβ-42 aggregation sites, potentially also inhibiting the re-aggregation of fibril fragments, thereby favoring the accumulation of likely toxic oligomeric species. This mechanism could explain the known adverse side effects of long-term L-dopa treatment in Parkinson’s disease, including irreversible motor dysfunction, such as dyskinesia^42^. Altered protein aggregation and the formation of toxic oligomeric species induced by L-Dopa may exacerbate neuronal damage and contribute to the observed motor complications, highlighting the delicate balance between L-Dopa therapeutic effects and the potential for exacerbating neurodegeneration^23, 43^. Our findings emphasize the complexity of L-Dopa interactions with pathological protein aggregates and the need to optimize treatment strategies to mitigate long-term side effects.

## Associated Content

### Author Information

#### Author Contribution

T.B. and S.C. prepared the α-Synuclein and Levodopa solutions. T.B. conducted the AFM measurements and data analysis for the α-Synuclein experiments. N.K. prepared the amyloid β-42 buffer solution and conducted the AFM measurements. P. N. N. conducted the AFM measurements on amyloid β-42 in cerebrospinal fluid and supervised the study. S.B. and D.T. performed the molecular dynamics simulations. T.B and P.N.N. wrote the manuscript. All authors discussed the results and commented on the manuscript. T. B and N. K contributed equally.

### Notes

The authors declare no competing financial interests

