## Supporting Information for "On Levodopa interactions with brain disease proteins at the nanoscale"

Talia Bergaglio[a,b], Nico Kummer[a], Shayon Bhattacharya[c], Damien Thompson[c], Silvia Campioni[d], Peter Niraj Nirmalraj[a,1]

[a]Transport at Nanoscale Interfaces Laboratory, Swiss Federal Laboratories for Materials Science and Technology, Dübendorf CH-8600, Switzerland; [b]Graduate School for Cellular and Biomedical Sciences, University of Bern, Bern CH-3012, Switzerland; [c]Department of Physics, Bernal Institute, University of Limerick, Limerick V94T9PX, Ireland. [d]Functional Materials Laboratory, Swiss Federal Laboratories for Materials Science and Technology, Dübendorf CH-8600, Switzerland

### Materials and Methods

#### Preparation of Wild-Type α-Synuclein, Aβ-42 and Levodopa solutions

Wild-type human α-Syn was obtained following the procedures outlined by Campioni et al[1] to prepare the α-Syn solutions, the lyophilized protein (~30 mg/mL) was first dissolved in 700 µL of PBS buffer (VWR). The pH was then adjusted to 7.4 using 1 M sodium hydroxide (NaOH). To filter the solution, the filter membrane of a 100 kDa NMWL centrifugal filter (Amicon Ultra-4 Centrifugal Filter Unit, Merk Millipore) was first hydrated with 4 mL of PBS buffer and centrifuged for 5 minutes at 3200g for three times. Next, the α-Syn solution (~700 µL) was added to the centrifugal filter and centrifuged for 20 minutes at 3200g to filter out any large α-Syn particles that had not fully dissolved. Lastly, to extract any remaining α-Syn from the bottom of the filter, 100 µL of PBS buffer was added to the filter and mixed, followed by centrifugation for 5 min at 3200g. A spectrophotometer (Implen Nanophotometer NP80 UV−vis) was used to determine the final concentration of α-Syn (ε280 = 5960 M−1 cm−1). The obtained α-Syn stock solution was further diluted with PBS buffer in order to reach a final concentration of 300 µM.

Aggregation of Aβ-42 in phosphate buffer salt solution (VWR) was performed according to the previously described protocol[2]. One entire vial (250 µg) of commercial Aβ-42 peptide (Sigma-Aldrich), was dissolved in 10% ammonium hydroxide (Sigma-Aldrich) at a concentration of 0.5 mg/mL by shaking at 400 rpm for 20 min. Aliquots containing 50 µg Aβ-42 were transferred into protein-low bind Eppendorf tubes and freeze-dried overnight. For each aggregation experiment, the freeze-dried pellet was dissolved in 100 µL 60 mM NaOH, (reaching a concentration of 0.5 mL or ~110 µM, confirmed by Implen NanoPhotometer at

280 nm with an extinction coefficient 1490 $M^{-1}cm^{-1}$and a molecular weight of: 4515 g/mol). The aggregation was initiated by diluting the sample with PBS to a final concentration of 5 μM and starting incubation at 37°C and 400 rpm shaking in an Eppendorf Thermomixer for 24h. To investigate the effect of L-Dopa on α-Syn and Aβ-42 aggregation dynamics, a stock solution was prepared by dissolving L-Dopa (Merk Millipore) in 1 mL PBS buffer (1.9719 mg/mL). For the samples incubated with L-Dopa, 250 μL of L-Dopa solution was added to 250 μL of α-Syn (300 μM), leading to a final concentration of L-Dopa of 100 μL. The α-Syn, with and without L-Dopa, was incubated at 37°C for 6 days under mechanical agitation at 300 rpm.

For Aβ-42 proteins in CSF (1.22 μg/L specified by vendor) we purchased the samples from Sigma Aldrich and the as-received samples were aliquoted and stored at -20°C until further use. The CAS number for this sample is ERMDA482IFCC.

**Atomic Force Microscopy**

Atomic force microscopy measurements on Aβ-42 aggregates in PBS incubated for 24 h were performed using a Bruker Dimension Icon instrument operated in tapping mode and equipped with SCOUT 150 HAR silicon AFM probes (gold reflective backside coating, force constant 18 N/m, resonant frequency: 150 kHz, NuNano). AFM measurements were conducted on air-dried α-Syn incubated without and with 100 μL L-dopa concentration on day 6 and deposited as a thin film on mica discs. The Aβ-42 solutions (~10 μL) were deposited on 5x5 mm Si wafers for 1 min and then rinsed with 1 mL ultrapure water, the excess water was dried under a gentle air stream. The raw AFM images were processed and analyzed using open source software Gwyddion 2.60. 2D leveling and scan line correction were applied, followed by measurements of the fibril height (Nuntreated = 316; NL-dopa = 277) and length (Nuntreated = 125; NL-dopa = 161). The size distribution of L-dopa particles observed in the treated α-Syn sample was calculated for a total of ~348 particles. Contour and persistence length measurements were performed on several AFM images recorded in different spots on the Si wafer using the matlab-based easyworm software[3]. Height profiles were extracted using the Bruker Nanoscope Analysis software. AFM measurements on Aβ-42 aggregates in CSF were performed using a multimode 8 Bruker instrument equipped with an E-scanner. For the AFM tip, a SCOUT 70 HAR silicon AFM tip was used in tapping mode (gold reflective backside coating, force constant 0.4 N/m, resonant frequency: 70 kHz, NuNano). AFM measurements were conducted on air-dried samples by first depositing 5 μL in separate experiments for both L-Dopa treated and untreated Aβ-42 aggregates in CSF medium on gold thin films followed by drying in air

(~5 h) and then placing the air-dried CSF samples on gold disks on top of the E-scanner for AFM imaging.

**Preparation of Models and Molecular Dynamics Simulations**

The atomically flat gold surface was modelled using a single-atom thick Au(111) slab of surface area 5 nm x 15 nm. By applying periodic boundary conditions (PBC) in the *xy*-plane, the infinite gold surface was modelled, and the Au atoms were held fixed with zero atomic charges. The Aβ-42 fibril structures were modelled as dodecamer (12-mer) in two different folds, one with two symmetric LS-shaped folds of hexamers packed laterally, obtained from the cryo–electron microscopy (cryo-EM) decamer (10-mer) structure of the Aβ-42 fibrils (PDB code 5OQV[4]), and the other with two symmetric double-horseshoe-shaped disease-relevant fold of hexamers packed laterally, and obtained from the solution NMR structure of a hexameric (6-mer) biological unit (PDB code 2NAO[5]). The initial structures of the pre-formed fibrils were oriented with the fibril axis parallel to the gold surface, with the two folds placed laterally on top of gold surface. The two fibril folds were placed at a minimum distance of all atoms of 5 Å above the gold substrate. 1000 molecules of L-Dopa were randomly added inside the simulation cell. To prevent excessive build-up of the L-dopa molecules on gold, the gold atoms were given zero Lennard-Jones parameters and the fibril models were given weak positional constraints on selected alpha carbon atoms to prevent protein dissociation from the gold nanosheet.

The fibril models were represented by the CHARMM36m[6] force field and solvated with CHARMM-modified TIP3P[6] water molecules. The topology and parameter of L-dopa was obtained from CHARMM General Force Field (CGenFF)[7, 8]. Molecular dynamics (MD) simulations were carried out using the Gromacs 2018.4[9, 10] package with a time step of 2 fs using the leapfrog integrator[11]. Bond lengths to hydrogen in protein were constrained using the LINCS[12] algorithm and water hydrogens were constrained using the SETTLE[13] algorithm. Background counterions were added to neutralise protein formal charges. Long-range electrostatics were treated by the Particle mesh Ewald[14] (PME) method. Protein and non-protein (gold, L-dopa, water and ions) were coupled separately to an external heat bath (298 K) with the coupling time constant of 1 ps using the velocity rescaling[15] method. The four systems (two fibril folds treated with L-dopa and untreated as control) were energy minimised, brought to room temperature over 100 ps and equilibrated for 1 ns in constant volume NVT ensemble followed by another 1 ns of constant pressure NPT equilibration with the reference pressure at 1 bar and a time constant of 4 ps using the Parrinello-Rahman[16, 17] barostat. The production runs were carried out for 100 ns for each of the four systems described above in the NPT ensemble. Structures were saved every 20 ps.

The trajectories were visualized and snapshots captured with Visual Molecular Dynamics (VMD)[18]. All analyses (main text Fig. 5 and Fig. S2 below) of root mean square fluctuation (RMSF), secondary structure and interaction energies analyses were performed using Gromacs tools. The fraction of native contacts was calculated using the definition from Best, Hummer and Eaton[19] implemented in the MDTraj[20] python library. The equation is:

$$\mathrm{Q(X)} = \frac{1}{N}\sum_{(i,j)}\frac{1}{1+\exp\left[\beta(r_{ij}(X)-\lambda r_{ij}^{0})\right]} \qquad \text{(eqn 1)}$$

where N is the set of all pairs of heavy atoms (i, j), and heavy atoms i and j are in contact if the distance between them is less than 5 Å and are separated by at least 3 residues. rij (X) is the distance between i and j in conformation X and $r^0_{ij}$ is the distance in the native. β is a smoothening parameter taken to be 5 $Å^{-1}$ and the λ is a factor that describes fluctuations when the contact is formed, taken to be 1.8 for the all-atom model. For more details on the methods and parameters, please see ref[19].

The conformational energy was estimated using the GBMV implicit solvent model (Generalized Born using Molecular Volume) implemented in the CHARMM (v40b2) program[21]. The energy was calculated after 200-step minimization of each MD snapshot using the GBMV II algorithm[22-24]. Other energy terms including bonded energy, van der Waals energy, electrostatic energy, and solvation energy were also calculated with the GB implicit solvent model. The block average method was used to estimate the mean values and standard deviations during the last 100 ns of dynamics, *i.e.*, 1000 statistically independent structures for each system.

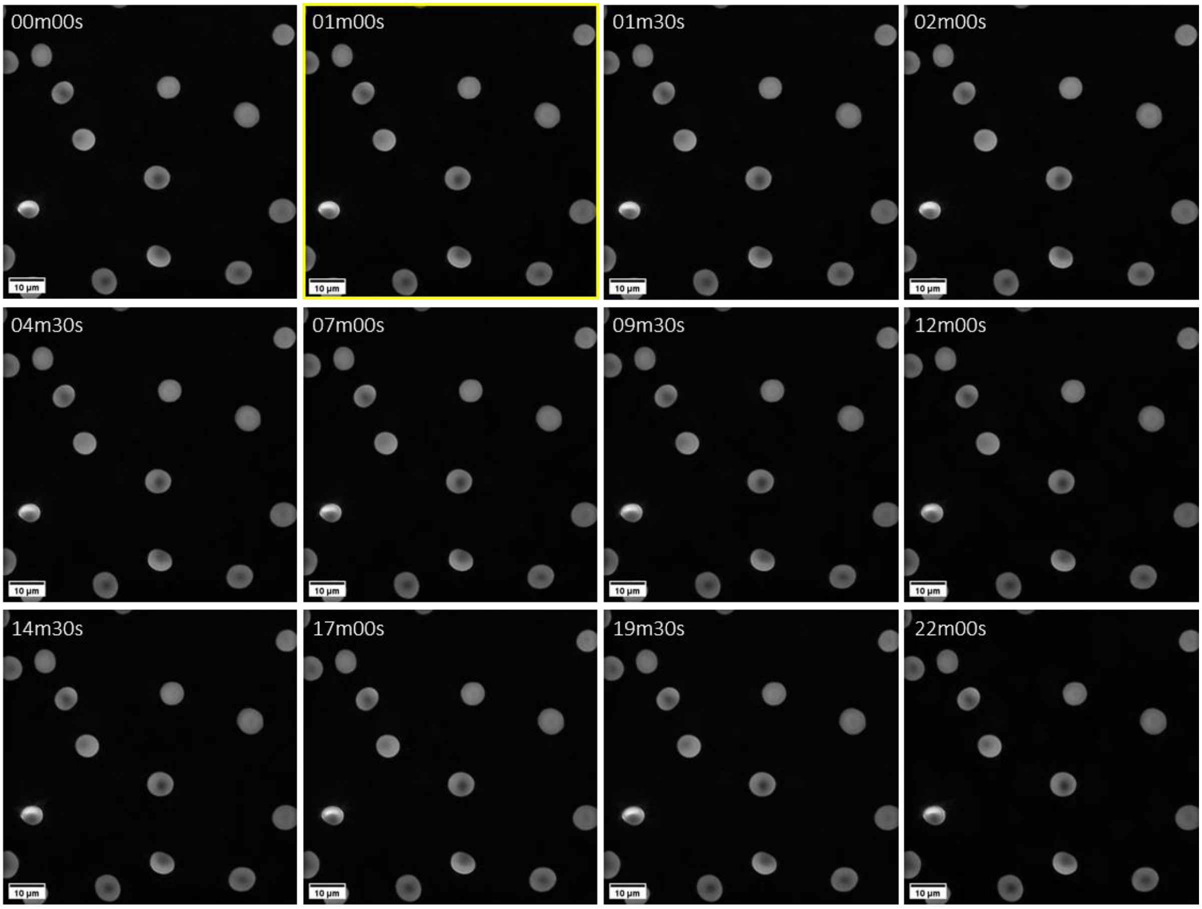


**Figure S1:**3D holo-tomographic imaging of red blood cells (RBCs) treated with 100 μM L-Dopa solution, during a 22-minute time-lapse. L-Dopa was added after 1 minute (yellow quadrant) from the start of the live cell imaging measurement. No structural alterations to RBC morphology were observed due to L-Dopa treatment.

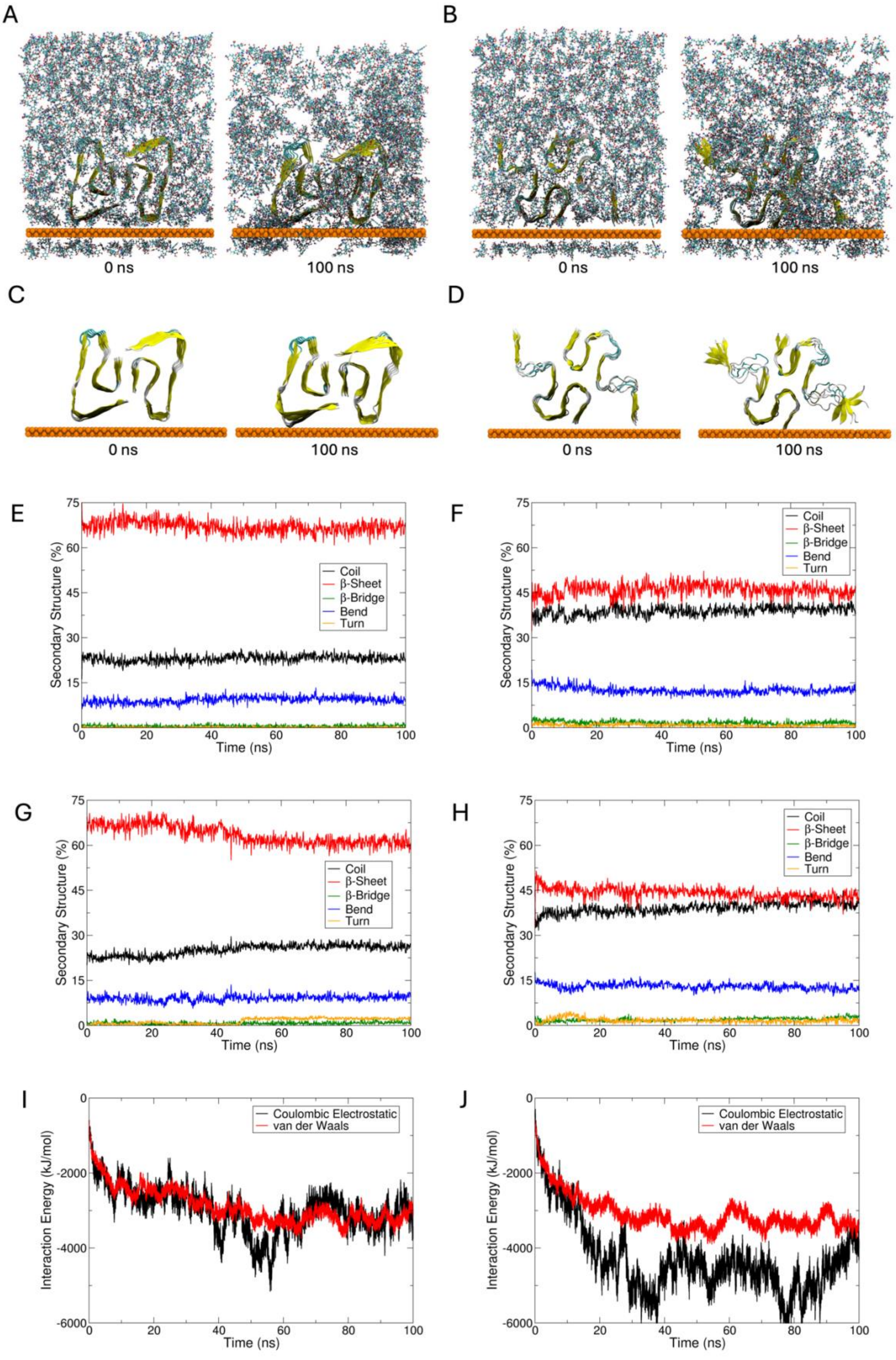

A
B
0 ns
100 ns
0 ns
100 ns
C
D
0 ns
100 ns
0 ns
100 ns
E
F
G
H
Coil
β-Sheet
β-Bridge
Bend
Turn
Secondary Structure (%)
Time (ns)
I
J
Coulombic Electrostatic
van der Waals
Interaction Energy (kJ/mol)
Time (ns)

**Figure S2:** The initial and final conformations of Aβ-42 fibrils in presence of L-dopa in the **(A)** LS-shaped fold, and **(B)** double-horseshoe-shaped fold from 100 ns MD simulations. The initial and final conformations of Aβ-42 fibrils in absence of L-dopa (control) in the **(C)** LS-shaped fold, and **(D)** double-horseshoe-shaped fold from 100 ns dynamics. Water molecules are omitted for clarity. The time evolution of percentage (%) of secondary structures of **(E)** LS-shaped fibril fold treated with L-dopa, **(F)** untreated LS-shaped fibril fold, **(G)** double horseshoe-shaped fibril fold treated with L-dopa, and **(H)** untreated LS-shaped fibril fold. Decomposition of interaction energies to their Coulombic electrostatic and van der Waals (vdW) components of L-dopa binding to (I) LS-shaped and (J) double-horseshoe-shaped fibril fold.
